# Age-dependent ribosomal DNA variations and their effect on cellular function in mammalian cells

**DOI:** 10.1101/2020.07.10.196840

**Authors:** Eriko Watada, Sihan Li, Yutaro Hori, Katsunori Fujiki, Katsuhiko Shirahige, Toshifumi Inada, Takehiko Kobayashi

## Abstract

The ribosomal RNA gene, which consists of tandem repetitive arrays (rDNA repeat), is one of the most unstable regions in the genome. The rDNA repeat in the budding yeast is known to become unstable as the cell ages. However, it is unclear how the rDNA repeat changes in ageing mammalian cells. Using quantitative analyses, we identified age-dependent alterations in rDNA copy number and levels of methylation in mice. The degree of methylation and copy number of rDNA from bone marrow cells of 2-year-old mice were increased by comparison to 4-week-old mice in two mouse strains, BALB/cA and C57BL/6. Moreover, the level of pre-rRNA transcripts was reduced in older BALB/cA mice. We also identified many sequence variations among the repeats with two mutations being unique to old mice. These sequences were conserved in budding yeast and equivalent mutations shortened the yeast chronological lifespan. Our findings suggest that rDNA is also fragile in mammalian cells and alterations within this region have a profound effect on cellular function.

**Author Summary:** The ribosomal RNA gene (rDNA) is one of the most unstable regions in the genome due to its tandem repetitive structure. rDNA copy number in the budding yeast increases and becomes unstable as the cell ages. It is speculated that the rDNA produces an “aging signal” inducing senescence and death. However, it is unclear how the rDNA repeat changes during the aging process in mammalian cells. In this study, we attempted to identify the age-dependent alteration of rDNA in mice. Using quantitative single cell analysis, we show that rDNA copy number increases in old mice bone marrow cells. By contrast, the level of ribosomal RNA production was reduced because of increased levels of DNA methylation that represses transcription. We also identified many sequence variations in the rDNA. Among them, three mutations were unique to old mice and two of them were found in the conserved region in budding yeast. We then established a yeast strain with the old mouse-specific mutations and found this shortened the lifespan of the cells. These findings suggest that rDNA is also fragile in mammalian cells and alteration to this region of the genome affects cellular senescence.

## Introduction

The genome, which comprises the complete set of genetic information in an organism, is sensitive to damage from environmental factors such as exposure to ultraviolet radiation. Damage to the genome is efficiently repaired by a highly organized repair system (1)(2). Nonetheless, some damage is not properly repaired leading to mutations, which may include rearrangements such as deletions and amplifications. In addition, mutations can also arise from errors introduced during DNA replication. These mutations accumulate during successive cell divisions to induce cellular senescence. However, the underlying mechanism linking accumulation of mutations to senescence is not well understood.

Damage to DNA tends to accumulate at fragile sites in the genome (3). In the budding yeast, *Saccharomyces cerevisiae*, the ribosomal RNA gene (rDNA) is known to be a fragile site that is related to cellular senescence (4). Eukaryotic rDNA is made up of repetitive tandem arrays, which in the case of the budding yeast comprises ∼150 rDNA copies located on chromosome XII. However, copies of these repeats are readily lost by homologous recombination. Because the cell requires a huge number of ribosomes, accounting for ∼60% of total cellular protein, a gene amplification system is needed to compensate for these losses. As a result, rDNA copy number frequently varies leading to an unstable genomic region (for review, see (5)). In terms of rDNA gene amplification in budding yeast, the replication fork blocking protein Fob1 works as a recombination inducer (6). Fob1 associates with the replication fork barrier (RFB) site, inhibiting the replication process and inducing a DNA double-strand break that triggers gene amplification/recombination (7)(8)(9).

Intriguingly, *fob1* mutants have a stable rDNA copy number, and lifespan is extended by ∼60% compared to the wild-type strain (10)(11). An important factor in suppressing rDNA copy-number change is Sir2, an NAD^+^-dependent protein deacetylase that is conserved across all kingdoms of life. Interestingly, *sir2* mutants of *S. cerevisiae* display increased unequal sister chromatid recombination, and the rDNA copy number frequently changes (7)(12). Moreover, the lifespan of the *sir2* mutant is shortened to approximately half that of the wild-type strain (13)(14). Taken together, these observations suggest that rDNA instability (i.e. frequent copy number alteration) is related to senescence (15).

In mammals, the rDNA structure is similar to that of yeast. However, the intergenic spacer sequence (IGS) in mammalian cells is larger than in yeast (Figure 1A) and is an unstable region of the genome (16). The connection between aging and rDNA has been suggested in several studies of tissues from dog, mouse and human (17)(18) (19)(20)(21). Werner syndrome is a human premature aging disease. The rDNA of cells derived from patients with Werner syndrome display an increased level of noncanonical arrangements (22). In the hematopoietic stem cells of mice, replication stress accumulates in the rDNA and cellular functional activity declines with age (23). However, there is still a paucity of observations how rDNA changes during senescence.

**Figure 1.**
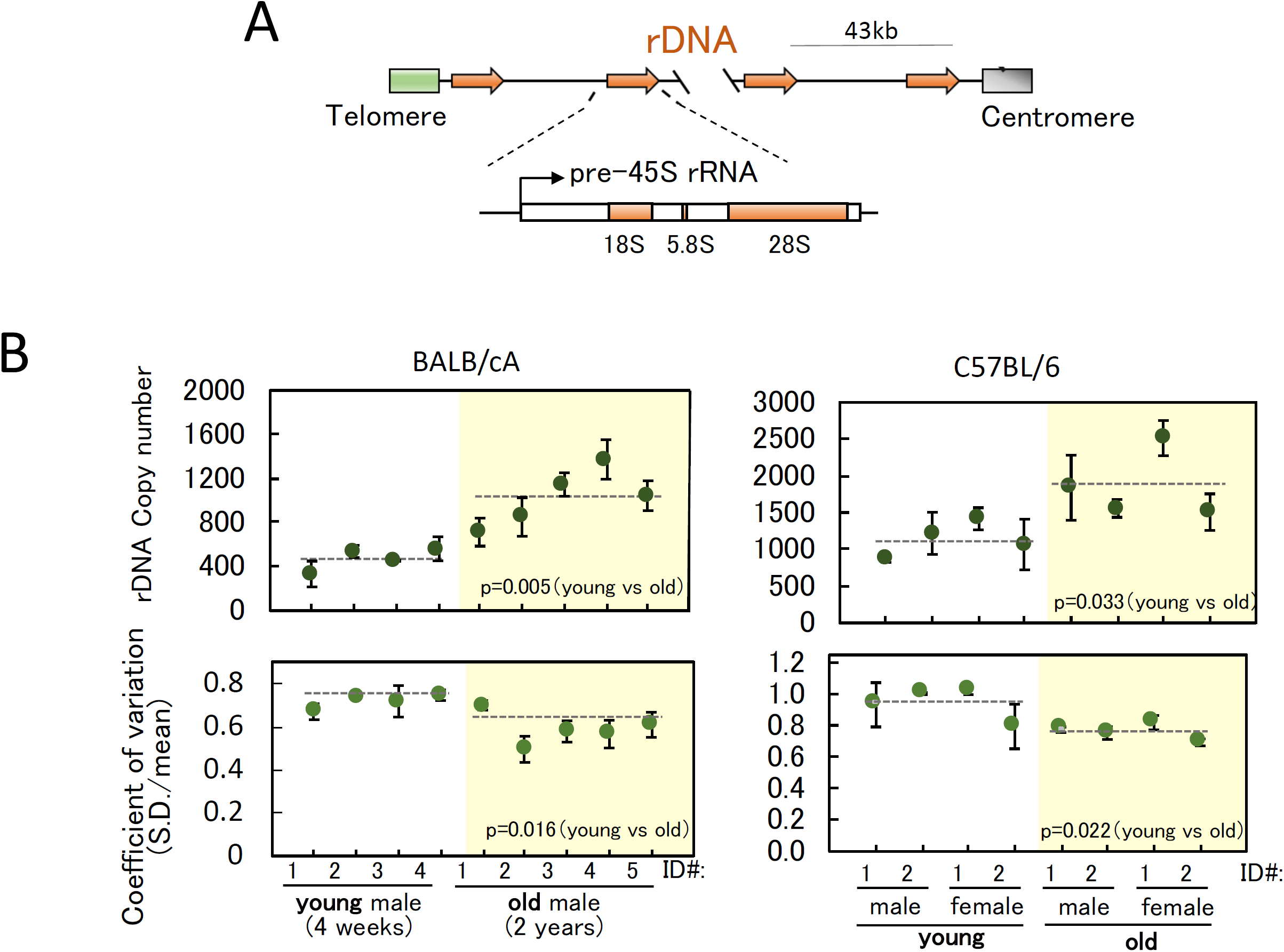
rDNA copy number and coefficient of variation in individual cells. **(A)** Structure of rDNA in mouse. One unit of rDNA is ∼43 kb, composed of the 45S pre-rRNA gene and intergenic spacer. 45S pre-rRNA is subsequently processed into three mature rRNAs (18S, 5.8S and 28S). (B) rDNA copy numbers were measured in young (4 week-old) and old (2 year-old) mice. (Upper) rDNA copy number in single cells were measured by qPCR and plotted. The copy number was determined using RPE1 as a standard (see Materials & Methods). The X-axis shows the identification numbers (ID#) of individual mice used to isolate the bone marrow cells. The dotted lines are the average values of the mice. The green dots are the average of two independent qPCR experiments from each mouse and the error bars indicate the range. (Lower) Plots of coefficient of variation (S.D. was divided by the average) for each mouse.

Here, we compared the genome of young and old mice, and identified differences in rDNA stability, methylation and transcription status. We also identified two mutations in rDNA that are specific to old mice. Moreover, these sequences are conserved in budding yeast rDNA. Interestingly, equivalent mutations in the budding yeast rDNA shortened their chronological lifespan. These findings suggest that the rDNA is also fragile in a mammalian cell and mutation of these sites affects cellular function.

## Results

### rDNA copy number is increased in older mice

Because the rDNA copy number readily changes, each cell may have a different copy number. Therefore, we initially measured the rDNA copy number in a single cell by quantitative real-time PCR (qPCR). In this strategy, we determined rDNA copy number of RPE1 (Human Retinal Pigment Epithelial cell) to obtain a standard curve by Droplet Digital PCR (ddPCR, BIORAD). In brief, a fixed amount of RPE1 DNA was digested into small fragments, diluted and fractionated into droplets. The dilution factor ensured that each droplet contains just one DNA fragment. Each droplet was then subjected to PCR and the number of positive droplets with an rDNA fragment counted. The ratio of the number of positive to negative droplets gives the absolute copy number of rDNA. Using this method, RPE1 cells were found to have 330 rDNA copies (See Materials & Methods for detail). The RPE1 DNA was then used as a control in determining the mouse rDNA copy number in a single cell by qPCR. Initially, we ensured the accuracy of the assay using one and two bone marrow cells, and one, two and four RPE1 cells to measure the rDNA copy number by qPCR. As anticipated, the rDNA copy number increased linearly with cell number (Figure S1).

Bone marrow cells were isolated from young (four-week old) and old (two-year old) BALB/cA and C57BL/6 mice. Specifically, four young and five old BALB/cA mice (males), and four young (two males, two females) and four old (two males and two females) C57BL/6 mice were tested. The cells were separated into a 96-well plate using a FACS machine and subjected to qPCR to determine the rDNA copy number. The results are shown in Figure 1B. We first noticed that the average rDNA copy numbers (dotted lines) are quite different in these two strains. In the young mice, they were 471 (BALB/cA) and 1,025 (C57BL/6) per cell, that is, C57BL/6 has more than double. The ratio (1,025/471) was 2.18. To confirm the difference, we also estimated rDNA copy number using publicly available whole genome sequencing data in NCBI. As shown in Figure S2, three mice data in each strain were analyzed and their average copy numbers were determined as 642 (BALB/cA) and 1,412 (C57BL/6) per cell. The ratio (1,412/642) was 2.20. Therefore, we think the difference of rDNA copy number in the two strains in our single cell analysis is reasonable and the analysis works well.

In terms of aging effect on the rDNA copy number, in both mouse strains, the average was increased in the older mice. We also calculated the coefficient of variation (S.D./mean) in individual cells, which indicates the rate of copy number variation in each mouse cell normalized by the average. The values obtained for the old mice were smaller than those for the young mice (see Discussion). These findings indicate that rDNA copy number increases in most old mice cells while the copy number variation decreases.

We also tested the copy number alteration in old mice by Southern blot analysis. In this assay, DNA was isolated from mouse bone marrow cells and double digested with BamHI/NdeI restriction endonucleases before being subjected to agarose gel electrophoresis (Figure 2). The probe for the Southern blot was designed to recognize the 28S rRNA gene in the 4 kb BamHI-NdeI restricted fragment. However, some of the rDNA copies had a second BamHI site (BamHI-2) in the 4 kb fragment (Figure 2A), resulting in the detection of two bands (Figure 2B, top). For BALB/cA, the upper bands (4 kb) appear stronger than the lower bands in the old mice, suggesting a relative loss of BamHI-2 sites within rDNA. To normalize the results, a single copy gene (SWI5) was also detected using a specific probe (Figure 2B, middle). The intensities of the bands were measured, and the values plotted (Figure. 2B, bottom). This analysis showed the intensity of the 4 kb BamHI - NdeI fragment for BALB/cA mice increased relative to the other fragments. Taken together, the data showed the rDNA copy number tended to increase with age although the difference was not as marked as in the qPCR analysis (See Discussion).

**Figure 2.**
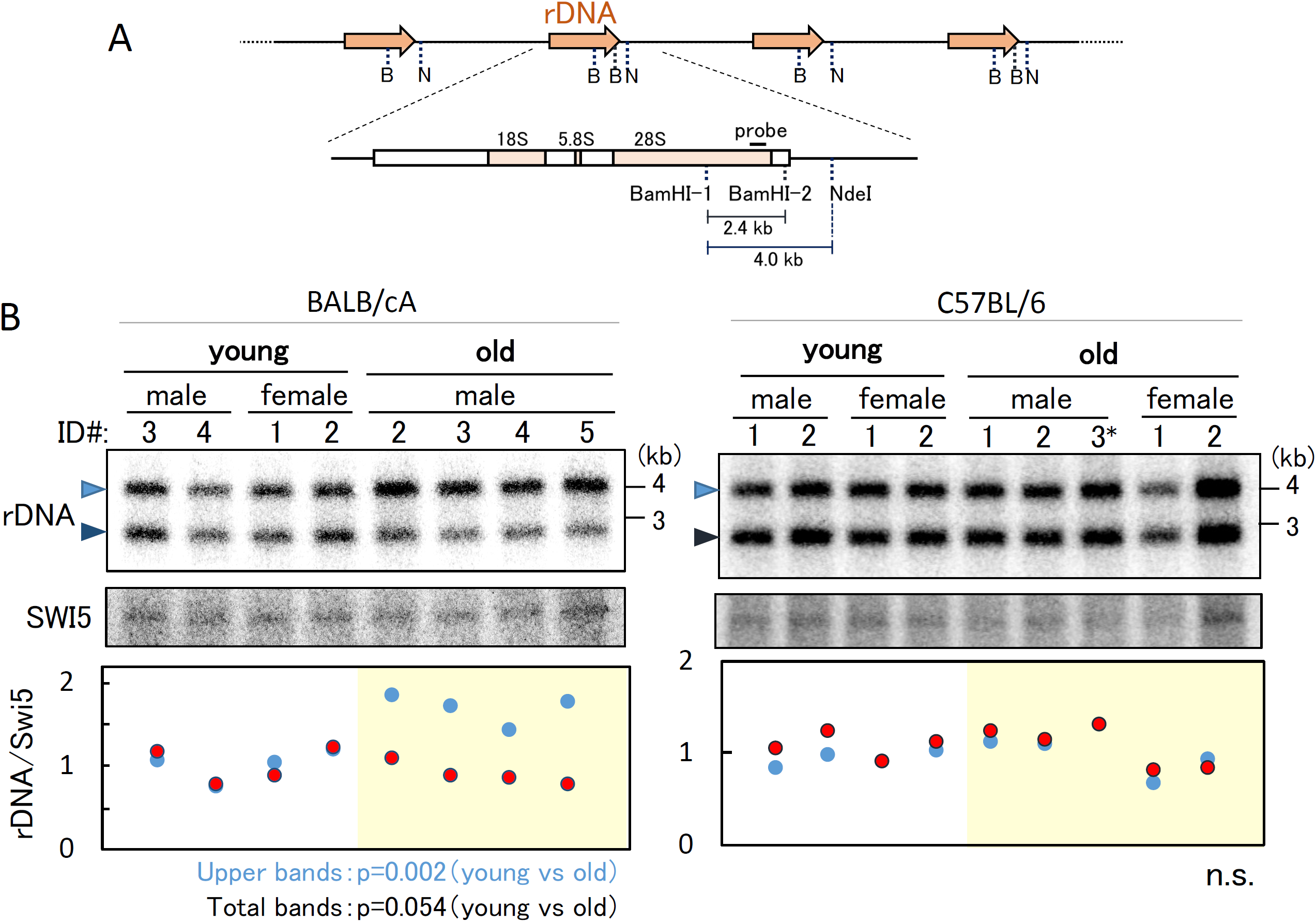
Detection of relative rDNA copy number in old and young mice. **(A)** Position of the probe for Southern blot analysis in (**BC**) and recognition sites for BamHI and NdeI are shown. (**BC**) Detection of relative rDNA copy number. (Top panel) Southern analysis for rDNA copy number. DNA was digested with BamHI and NdeI. Upper bands (4 kb) come from rDNA units without BamHI-2 site and lower bands (2.4 kb) from rDNA units with BamHI-2 site. (Middle panel) Detection of SWI5 gene as a loading control. A single copy gene SWI5 was detected on the same filter used in the upper panel. (Bottom panel) Relative amount of rDNA copy number. The band intensities of rDNA were normalized by those of SWI5 and the values are relative to the average of rDNA values in the four young mice. The blue dots show the results from the upper band intensities of rDNA and the red dots are the results from the lower bands. ID# is the identification number of individual mice that were used to isolate the bone marrow cells (Figure 1). p values are shown at the bottom of the panel. n.s. is “not significant”.

### rDNA transcription levels are decreased in the older mice

The previous qPCR and Southern analysis showed the copy number of the rDNA tended to increase in older mice. We therefore speculated that the increased copy number of rDNA might result in an elevated level of rDNA transcripts (rRNA). To test this hypothesis, RNA was isolated using cells derived from young and old mice and the level of 28S rRNA measured by RT qPCR. The values were normalized against the transcripts of three housekeeping genes, Actb (actin, beta), B2M (beta-2 microglobulin), and GAPDH (glyceraldehyde-3-phosphate dehydrogenase). The results are shown in Figure 3B. Although there was a tendency for the young mice cells to have more 28S rRNA, the difference was not significant except for the results normalized against B2M. It is possible that the housekeeping genes are also affected by age. In addition, most of the 28S rRNA are thought to be included in the ribosomes that abundantly accumulate in the cell. Therefore, we measured newly synthesized pre-matured 45S rRNA using a probe that recognizes the promoter region and then calculated the ratio of matured to pre-matured rRNA. As shown in Figure 3C, in BALB/c, the newly synthesized rRNA ratio was reduced in the old mice. However, this difference was not as obvious in the C57BL/6 mice.

**Figure 3.**
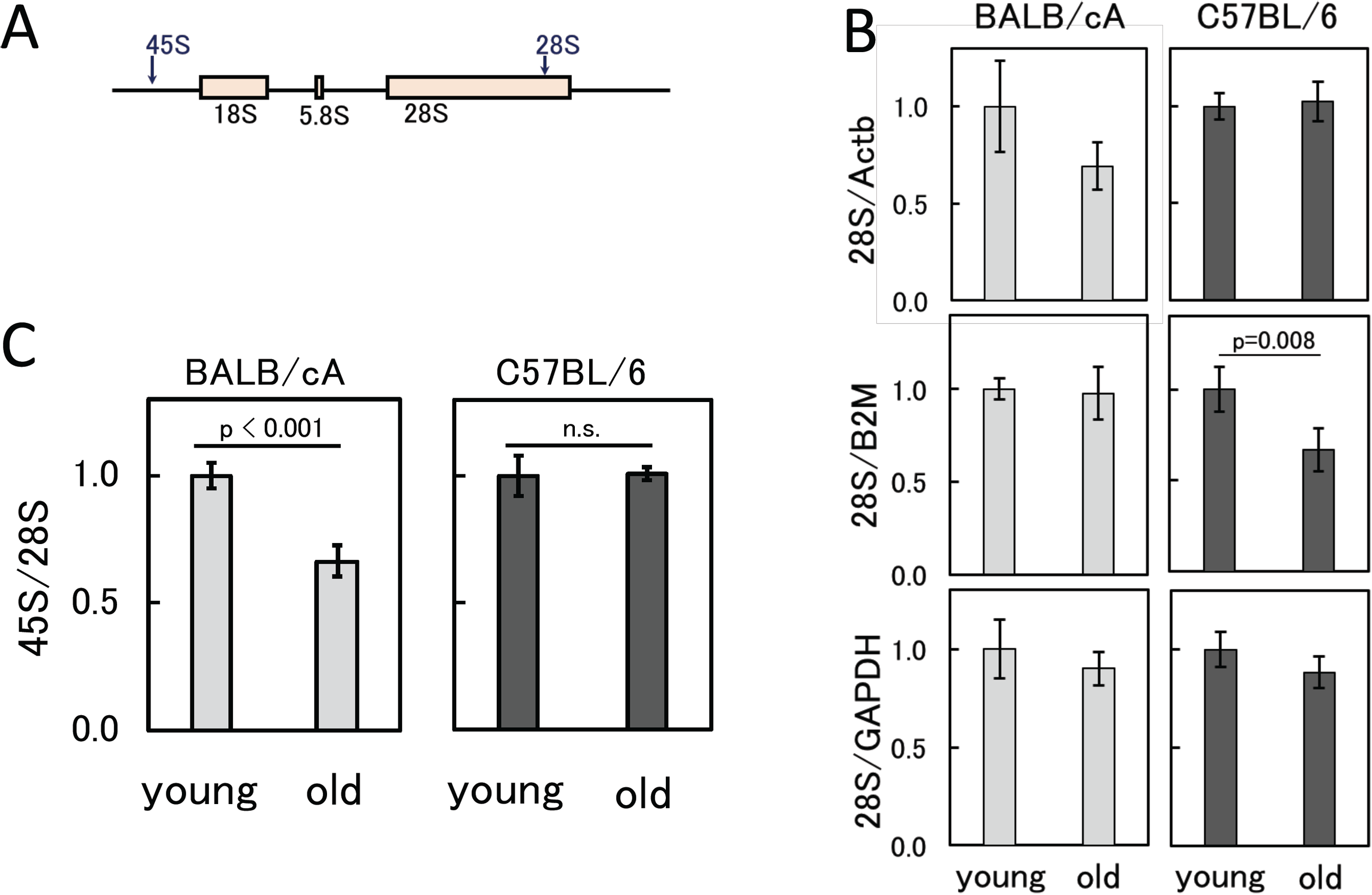
rDNA transcripts in old and young bone marrow cells. (**A**) Positions of the primer sets for qPCR to measure rDNA transcripts (pre 45S and 28S rRNA). (**B**) Amount of 28S rRNA normalized by transcripts of the three housekeeping genes (GAPDH, B2M and Actb). The values are the average of four independent experiments and the error bars are S.D. The values are relative to those of young cells. p value is shown in case it is significant (p<0.05). (**C**) Ratio of the pre 45S rRNA to 28S rRNA. The values are the average of four independent experiments and the error bars are S.D. The values are relative to those of young cells. p value is shown. “n.s.” is “not significant”.

Transcription inactivation of rRNA gene in C57BL/6 mice was confirmed using the psoralen crosslinking method (24). Psoralen intercalates into non-nucleosomal rDNA copies that are actively transcribed more efficiently than those that are transcriptionally inactive. Therefore, using this method, we can estimate the proportion of active rDNA copies. Cells are treated with psolaren, UV crosslinked and the DNA isolated. After digestion with AflIII the DNA was subjected to agarose gel electrophoresis. The results are shown in Figure 4. The upper and lower bands correspond to transcribed (active) and non-transcribed (inactive) rDNA copies, respectively (Figure 4B). Band intensities were measured, and the values plotted (Figure 4C). The ratio of active to non-active rDNA was less in the old cells than in the young cells. These findings suggest that rDNA transcription is reduced in the older mice.

**Figure 4.**
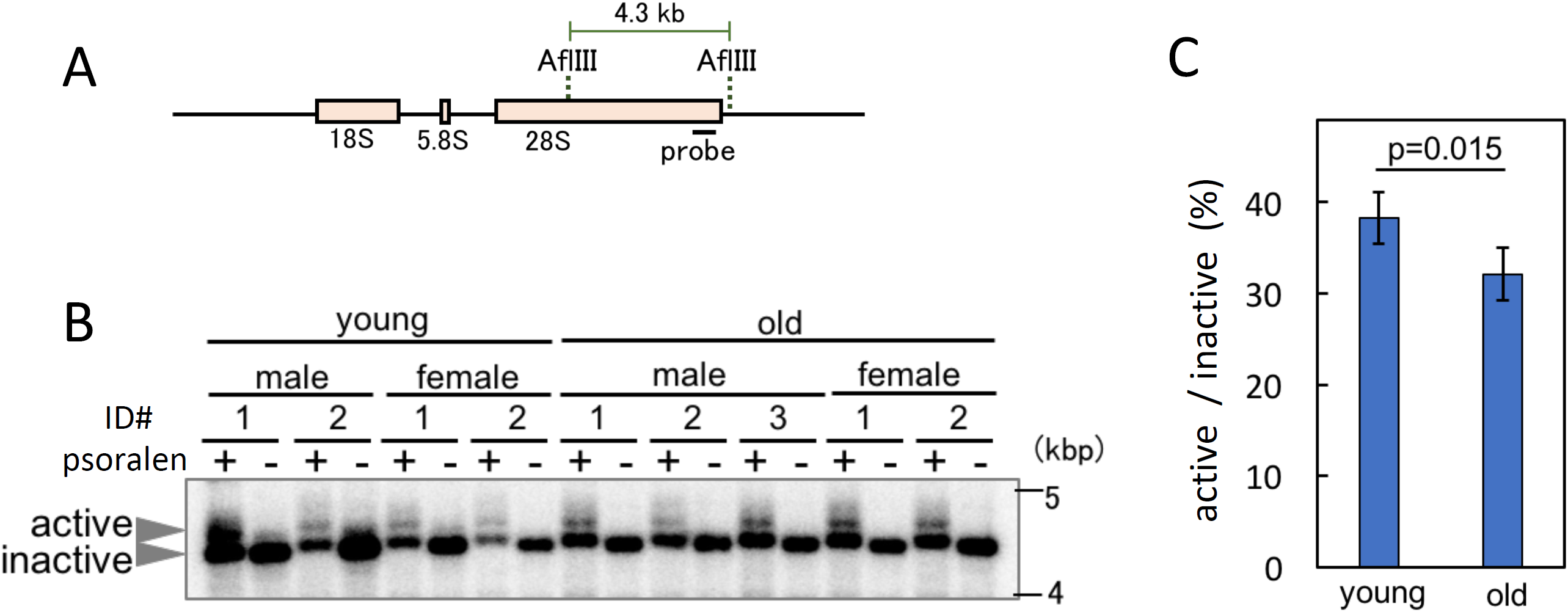
Ratio of active and inactive rDNA copies in old and young cells. Psoralen crosslinked rDNA was digested with AflIII and band retardation assessed after electrophoresis. (**A**) Recognition sites of AflIII and position of the probe for (**B**). (**B**) Southern blot analysis to detect the psoralen crosslinked rDNA by mobility retardation in young and old mice. Transcription “active” rDNA efficiently intercalates psoralen, which retards migration during gel electrophoresis. ID # is the identification number of the mice (same as Figure 2B). (**C**) Ratio of active to inactive rDNA copies. Band intensities of (**B**) were measured and the ratio of active to inactive rDNA calculated. The error bars are S.D. The p value is shown.

### rDNA is more highly methylated in the older mice

Transcription is known to be affected by DNA methylation (25). Recently, it was reported that the methylation rate of rDNA increases in an age-dependent manner in both mouse and human (26). Therefore, we speculated that increased methylation of rDNA might reduce the transcription level in older mice. To test this hypothesis, DNA from the old and young mice was digested using a methylation sensitive enzyme SacII and the restriction pattern analyzed (27). As shown in Figure 5B, in the absence of SacII, two bands (4.0 and 2.4 kb, highlighted by arrowheads) were observed after BamHI-NdeI digestion (refer to Figure 2B). However, after SacII digestion most of these bands disappeared in the young mice. By contrast, the same analysis of DNA from old mice showed faint bands were still detectable (Figure 5B). The signal intensities of undigested and digested bands were measured, and the ratios calculated. As a loading control, a single gene SWI5 was also detected. The values of signal intensity were then plotted (Figure 5B, lower panel). The ratios of methylation in the old mice were increased except for the #2* mouse. The same assay was performed in C57BL/6 strain and similar results were obtained (Figure 5C). These results confirmed that rDNA in the old mice is more methylated than in the young mice. Taken together, our findings suggest DNA methylation causes the reduced level of transcription of rDNA.

**Figure 5.**
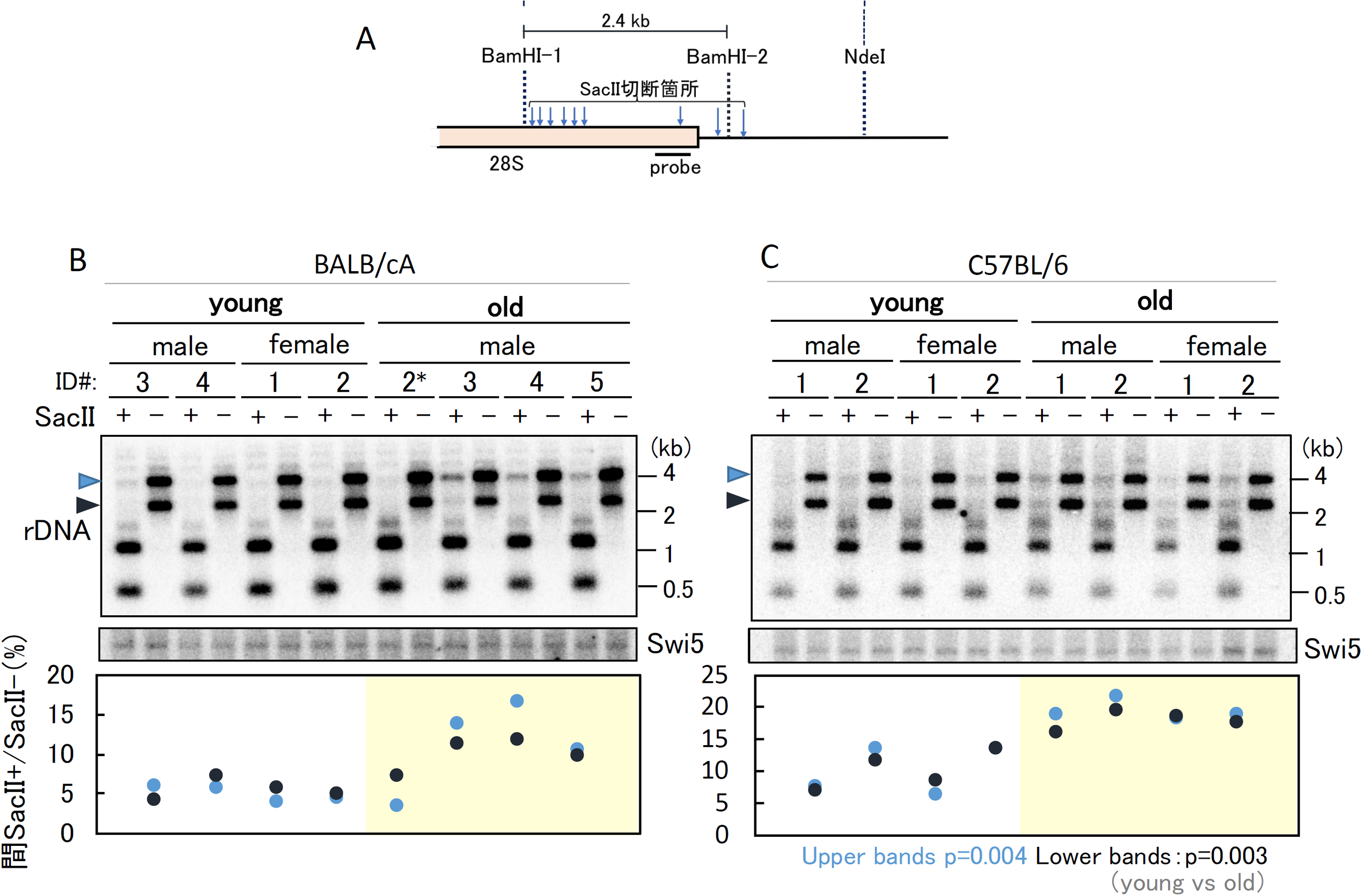
rDNA methylation in old and young bone marrow cells. rDNA methylation was detected by digestion with a methylation sensitive enzyme SacII. (**A**) Position of the probe for Southern blot analysis in (**BC**) and recognition sites for BamHI and SacII. (**BC**) Ratio of methylated rDNA copies in young and old mice. (Top panel) Southern analysis to detect the ratio of undigested band by SacII. The positions of undigested bands are indicated by arrowheads on the left-hand side of the panels. (Middle panel) Detection of the SWI5 gene as a loading control. A single copy gene SWI5 was detected on the same filter used in the upper panel. (Bottom panel) Analysis of rDNA that failed to digest with SacII. The signal intensities of the undigested rDNA (SacII+) and non-digested (SacII-) bands were measured and the ratios plotted. The black dots show the results for the lower bands (black arrowhead, in the Top panel) and the blue dots are for the upper bands (blue arrowhead in the Top panel). ID# is the identification number of individual mice that were used to isolate the bone marrow cells (same as Figure 2). p values are calculated from the average of young and old mice.

### There is sequence variation in rDNA of young and old mice

Finally, we determined the rDNA sequence in the young and old mice. Bone marrow cells, including hematopoietic stem cells that produce leukocytes, erythrocytes and platelets, are known to divide frequently. Thus, we speculated that mutations in the older mice cells accumulate and affect the function of the ribosome causing aging phenomena, such as slow growth and reduced viability. DNA from young and old mice was isolated and the 18S, 5.8S and 28S genes PCR amplified for analysis by deep sequencing. All of the reads were aligned and compared with the mouse reference sequences (28) to identify mutation sites. The results are shown in Figure 6A-C. The “mutation rate” is the ratio of mutations identified in the sequences to the total reads. Thus, a “mutation rate of 1 (100%)” means the sequence is different from the reference sequence. If the mutation rate is 0.5 (50%), half of the rDNA copies display a variation at that site. As a control, we also analyzed a housekeeping gene ATP5b (ATP synthase gene) (Figure 6D).

**Figure 6.**
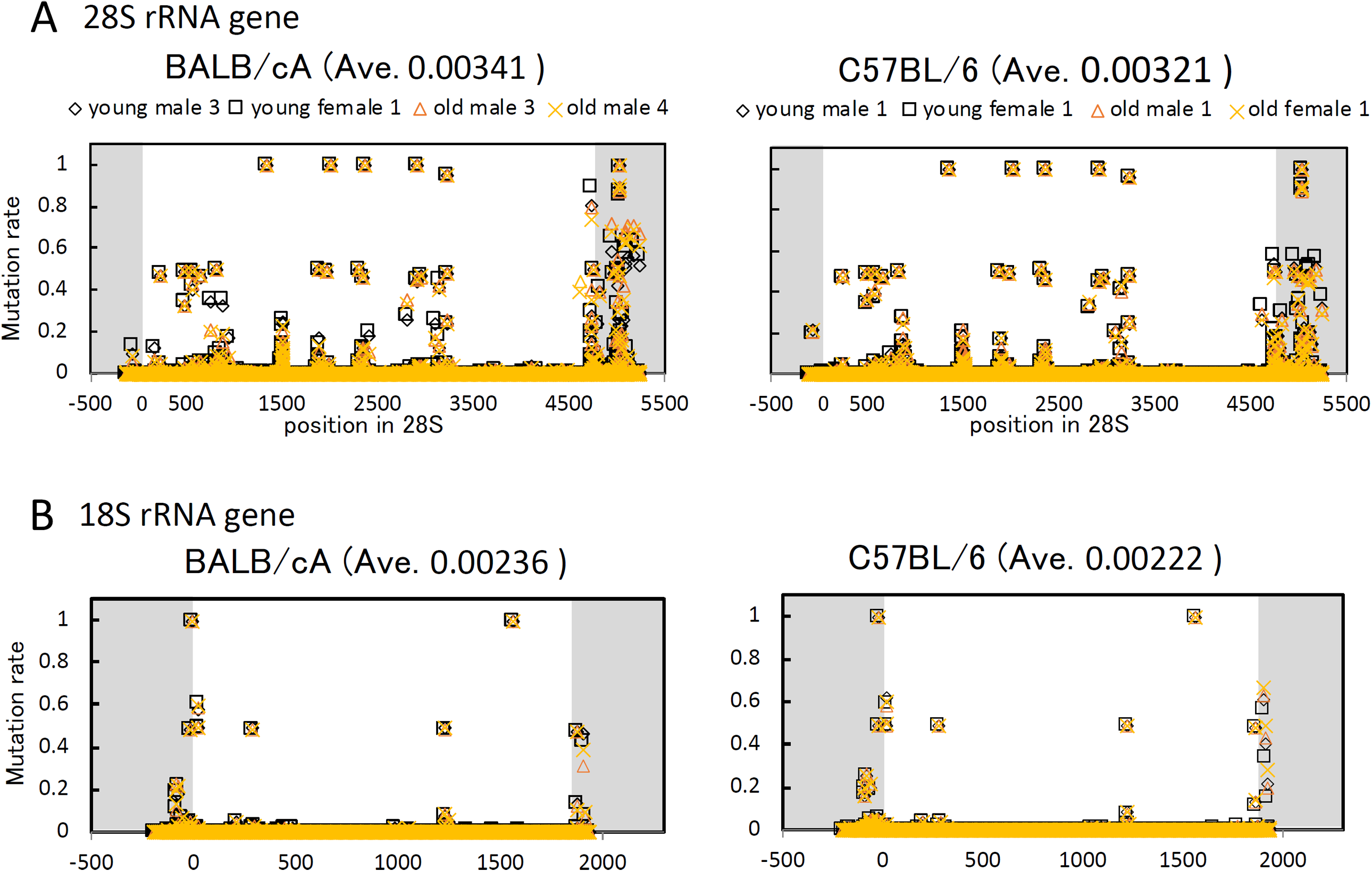
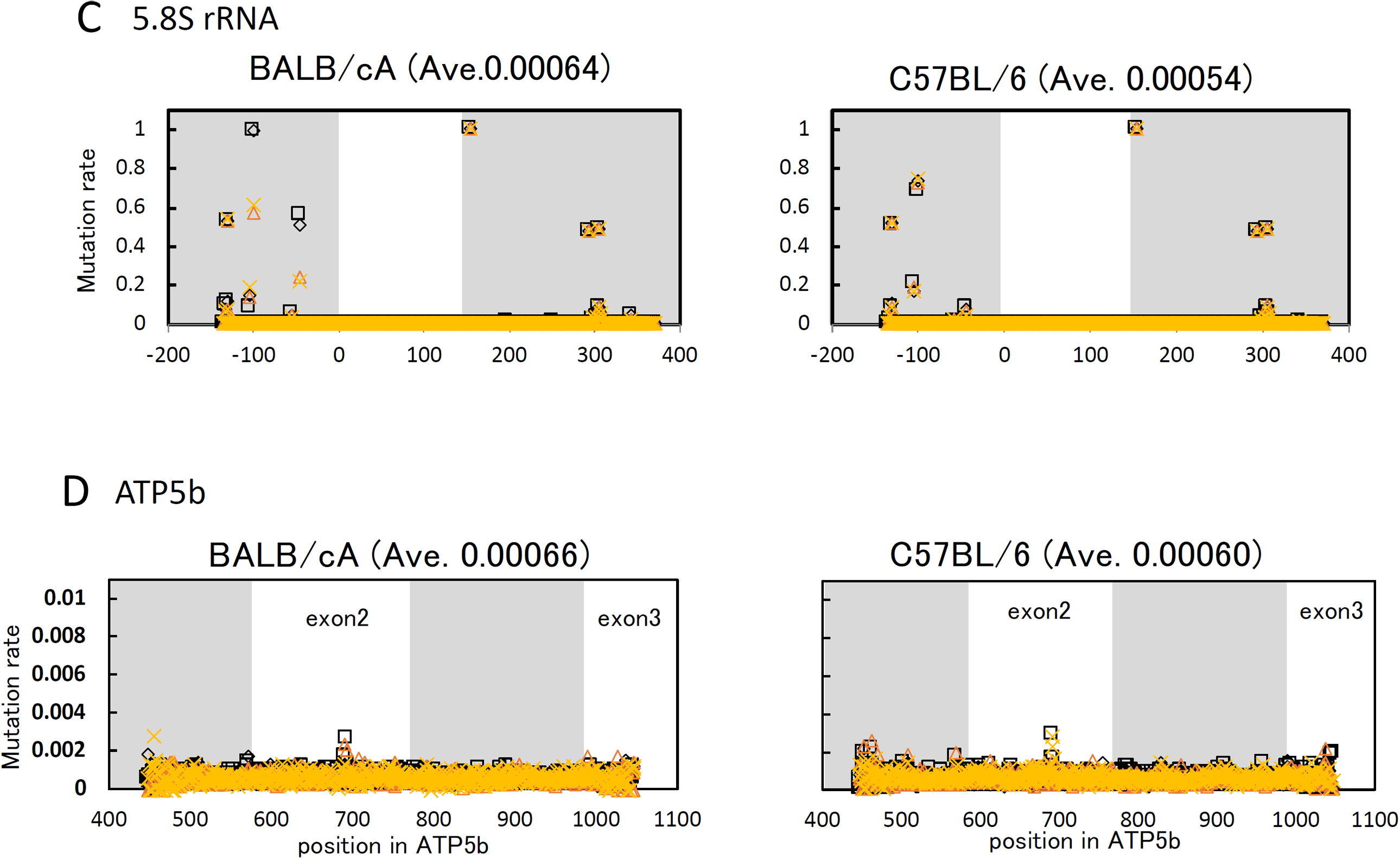
rDNA sequence variation in young and old mice. The rDNA sequences from two old and two young mice were determined and compared to the reference sequence. Mutation rates were then calculated at each nucleotide position. (**A**) 28S rRNA gene, (**B**) 18S rRNA gene (**C**) 5.8S rRNA gene and (**D**) ATP5b gene. Non-coding regions are not shadowed. Mutation rate is the difference from the reference sequence. Ave. is the average mutation rate.

As shown in Figure 6, the overlapping black and yellow marks indicate the mutation rates in the young and old mice cells were similar. The average mutation rates in both young and old mice cells were similar (Figure S3 and S4E). Thus, any age-dependent alteration of rDNA sequence was not immediately apparent. Nonetheless, the average mutation rate of 28S rDNA (BA:0.00341, BL:0.00321) was higher than that of 18S rDNA (0.00236, 0.00222) and much higher than ATP5b and 5.8S (0.00054∼0.00066). Indeed, sequence variation among copies of 28S rDNA has been reported previously (29). All of the high rate variations in 28S and 18S rDNA were found in DNA from both young and old mice (Figure 6).

For the purpose of identifying old mice-specific mutations, we searched for variations with a mutation rate of >0.0028 (0.28%), which was equivalent to the maximum value for the control gene (ATP5b). The threshold value is the maximum apparent artificial mutation rate caused by PCR amplification or other errors. Within this range, we identified three old mice-specific mutations in the old BALB/cA strain (Table 1). By contrast, no old mice-specific mutations were identified in the C57BL/6 strain. Indeed, no old mice-specific variations were found after increasing the number of mice that were sequenced (Figure S4).

**Table 1.** The positions of old mouse specific mutations in BALB/cA. The values refer to the mutation rates. Position 3094 is not conserved in yeast rDNA.

| positions<br>of mouse<br>28S | young |  | young<br>ave. | old |  | old ave. | old –<br>young | positions<br>of yeast<br>25S |
| --- | --- | --- | --- | --- | --- | --- | --- | --- |
|  | Male #3 | female #1 |  | male #3 | male #4 |  |  |  |
| 4614 | 0.000 | 0.001 | 0.001 | 0.441 | 0.396 | 0.419 | 0.418 | 3295 |
| 3291 | 0.001 | 0.001 | 0.001 | 0.039 | 0.032 | 0.035 | 0.034 | 2131 |
| 3094 | 0.036 | 0.038 | 0.037 | 0.009 | 0.007 | 0.008 | - 0.029 | – |

Accuracy of the sequencing data was verified by analyzing variation of the BamHI recognition sequence that was detected in Figure 2 and Figure 5. The anticipated variation in the sequencing data corresponding to the BamHI site (GGATCC) in both mouse strains was observed together with the changes seen in the old BALB/cA mice (0.25 to 0.685) (Table S1). Thus, the sequencing data correlate with the Southern analysis in which the intensities of the upper bands increased in the old BALB/cA mice (Figure 2B).

### The old mouse-specific mutations of rDNA affect yeast ribosomal function

To analyze the relationship between rDNA variation and function, we summed up the mutation rates in 20 bp windows and plotted the values (Figure 7). In the graph, several variations, or “hotspots”, were identified over the 28S rDNA. Interestingly, most of the hotspots (highlighted in yellow) were located in the non-conserved regions between mouse and budding yeast rDNA (red line, top). These observations suggest that most of the variations are present in the non-functional region of the 28S rRNA gene.

**Figure 7.**
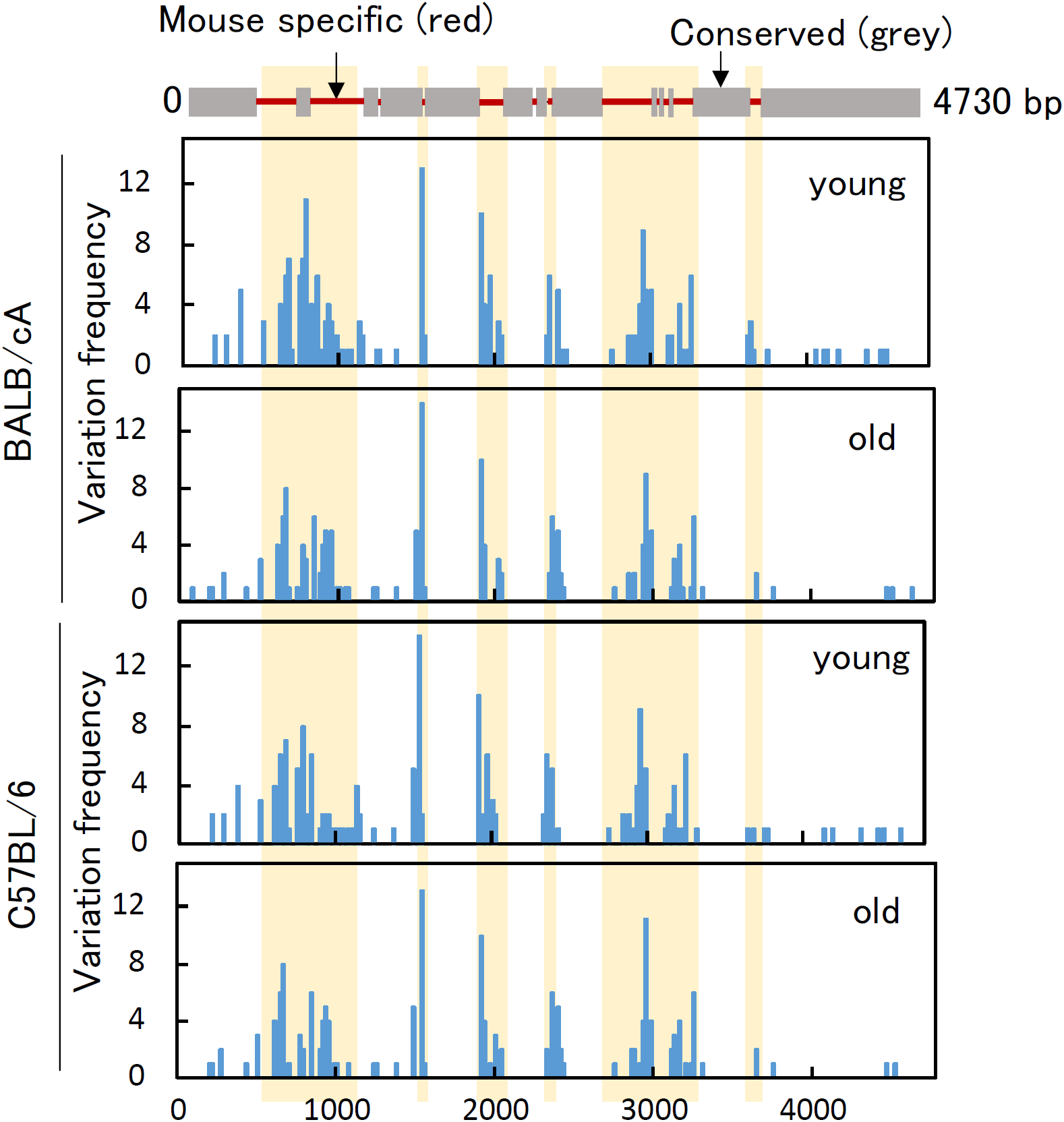
Hotspot of variation in the 28S rDNA. Mapping of hotspots of variation in mouse 28S. (Upper panel) Alignment of mouse 28S and budding yeast 25S rRNA genes. The gray boxes are conserved regions in the two species and the red lines are mouse unique regions. (Lower panel) Variation frequency in the 28S rRNA gene. The mutation rates were summed (more than 0.0028 was picked up, Figure 6A) every 20 bp and then plotted. Data from two mice were used in each graph. The yellow boxes represent mouse specific non-conserved regions.

We also mapped the positions of the three old mouse-specific mutations identified in the BALB/cA mice to yeast 25S rDNA. Interestingly, two sites (3291 and 4614) were plotted in the conserved region between mouse and yeast, suggesting they might be located in the functional domains in the rRNA. One approach to study the consequence of these mutations is to examine their impact in yeast. Thus, we generated budding yeast strains carrying the corresponding mutations in the 25S rDNA. For specific expression of the mutated rDNA, we used a yeast strain without rDNA in the chromosome (*rdnΔΔ* strain) (30). The strain initially carried a helper rDNA plasmid, which was then shuffled with plasmids containing mutations in the 25S region. The plasmid-borne mutated rDNA thus became the sole source of rRNA. Strains with either plasmid-derived wild-type rDNA, A2131G (mouse A3291G), or A3295G (mouse A4614G) mutated rDNA showed comparable cell growth in both solid and liquid medium. To test the relationship between these mutations and senescence, we measured the chronological lifespan by calculating survival rates every two days after the cells entered the stationary phase. As shown in Figure 8, one of the mutations (A3295G) lowered the proportion of surviving cells at all time points from day 5 onwards, indicating a shortened chronological lifespan. By contrast, another mutant (A2131G) showed similar survival rates to that of the wild-type yeast until day 15, but then the rate dropped on day 17. These observations suggest that although both mutations identified in the old mouse rDNA support cell growth in yeast, they may be harmful during chronological aging, particularly A3295G (mouse A4614G).

**Figure 8.**
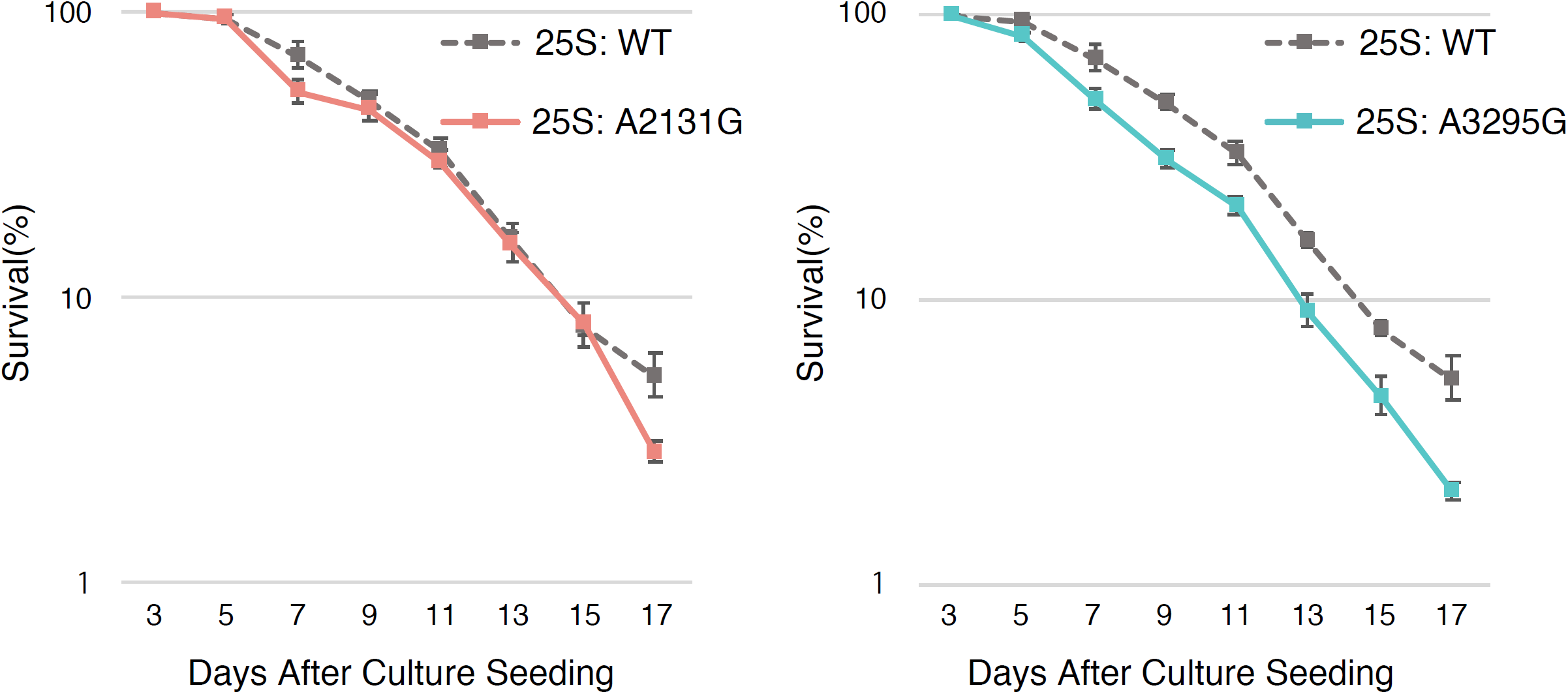
Chronological lifespan in budding yeast with mutated rDNA. Two old mouse-specific mutations in the 28S rRNA gene were introduced into the budding yeast 25S rRNA gene and the chronological lifespans of the yeast measured. Lifespans in the yeast strains with A2131G (left panel) and A3295G mutations in the 25S rRNA gene. The values are the average of nine experiments.

## Discussion

The rDNA has the following unique features that make it possible to monitor age-dependent alterations in the genome. Firstly, because rDNA is a highly repetitive and recombinogenic region it is easy to assess instability by monitoring alterations in copy number. Secondly, as approximately half of the rDNA copies are not transcribed (24)(31), these repetitive non-transcribed regions are targets for both methylation (32) and mutation (33). Indeed, our analyses detected alterations in copy number and methylation level in old mice, as well as putative old mouse-specific mutations.

In terms of rDNA copy number alteration observed in old mice, the results from literature reports are contradictory (18). Copy number alteration itself is commonly observed by many researchers, but in some reports the number goes up and in others it goes down. Moreover, copy number alteration has also been observed in tissues (18). Some of these discrepancies may arise from problems related to hybridization during Southern blot analysis. The repetitive nature of the DNA combined with the high level of bound proteins from the nucleolus may affect the detection efficiency. Indeed, our results showed that although the rDNA copy number in old mice increased as detected by single cell analysis by qPCR, this increase was not obvious by Southern blot analysis in either of the two mouse strains (Figure 1 and Figure 2). For budding yeast, rDNA copies in the old cells dramatically increases (∼10 times) as extra chromosomal rDNA circles (ERC) and their presence is a big burden on the cells because ERCs consume factors that are required for chromosome maintenance (34)(35). Therefore, the copious amount of ERC is thought to be a passive accelerator of cellular aging. In the case of mammals, this age-dependent increase of rDNA copies is not as dramatic (< 2 times, Figure 1). As such, the extra rDNA copies in mammals may not in itself reduce lifespan.

In terms of genome instability, it may be possible to connect age-dependent changes in rDNA to the aging process. To address this issue, we previously established a strain of *S. cerevisiae* with reduced replication initiation activity only in the rDNA (36). Because ERC cannot replicate, there is no ERC accumulation. However, the lifespan of the strain was shortened, and rDNA stability was reduced in the strain. We speculated that extended travel of DNA polymerase, due to reduced replication initiation, induces DNA replication stress, such as fork arrest and damage, leading to genome instability. These findings suggested rDNA instability and/or damage itself is an aging signal that shortens lifespan (5). From this viewpoint, yeast and mammalian rDNA may play similar roles in terms of aging by acting as a large fragile site for disseminating an aging signal (5). Indeed, the replication fork blocking activity that causes rDNA recombination in the budding yeast is also present in mammalian rDNA (for a review see (37)(38)). A similar fork arrest induces rDNA instability to promote senescence by distributing the aging signal. Further studies are required to investigate this hypothesis.

In the single cell analysis, we found that the copy number of rDNA increased and the variation decreased in older cells. As far as we are aware, there is no previous report showing alteration of rDNA copy number at the single cell level. One possible reason to explain the reduced variation phenotype in the old cells is that the number of stem cells for bone marrow goes down with age. Indeed, it is known that the number of the hematopoietic stem cells in the bone marrow gradually decreases during the process of aging (39). As bone marrow cells are produced from the stem cells, the variation of rDNA copies is reduced. The relationship between rDNA methylation and senescence has been discussed in previous reports (25)(26). The present results are consistent with these previous studies in showing that rDNA is more highly methylated in older mice (Figure 5). DNA methylation is known to repress transcriptional activity (25). Indeed, the ratio of 45S to 28S transcripts reduced in the old BALB/cA mice. The underlying reason for the age-dependent increase in methylation has not been elucidated. However, repetitive non-coding elements, such as retrotransposons, are known targets for DNA methylation enzymes (32). In addition, rDNA is subject to DNA damage and has a high GC content, which are known to be related to age-dependent methylation (40)(41). Hence, a similar mechanism may recognize the repetitive rDNA as a target for methylation. Moreover, in terms of the relationship between reduced rDNA transcription and increased copy number in old cells, one possible explanation is that cells can compensate for lowering the production of rRNA by elevated copies of rDNA to enable them to survive. As a result, the rDNA copy number in the old mice is more than in the young mice.

We anticipated more mutations in the older mice because there are many untranscribed non-canonical rDNA copies (22) and hematopoietic stem cells are subject to DNA replication stress (23). The untranscribed copies can accumulate mutations and replication stress increases DNA damage. However, the mutation rate in old mice was similar to that in young mice (Figure S3 and S4E). Therefore, cells should have an effective repair system and/or mechanism to avoid mutation accumulation such as gene conversion for homogenization (33). In this study, we identified three such mutations in the old mice. Although these mutations were present only in the old mice, it is not known whether they occurred during the aging process. Moreover, it is not known whether the rDNA copies with the mutation are actually transcribed or not. Thus, these mutations may not be related to senescence in the mice. Nonetheless, we found that two equivalent mutations in the budding yeast permitted cell growth, but one of the mutations (A3295G) apparently shortened the chronological lifespan. These findings indicate that the mutated rDNA, when present as the only source of rRNA, is transcribed and can support the essential functions of the ribosome, but viability during aging is negatively impacted, at least in yeast. Therefore, one could infer that if such harmful mutations accumulate in the rDNA repeats during the course of successive cell divisions, they may cause defects in the ribosomal and cellular functions to induce senescence.

In this study, we used two mice strains, BALB/cA and C57BL/6, for the analyses and they showed slightly different results. The rDNA copy number in C57BL/6 is twice as large as that in BALB/cA. Age-dependent alterations in the copy number, transcription and methylation levels were more prominent in BALB/cA. The mutation rate in BALB/cA was also higher than that in C57BL/6 and we were only able to identify specific mutations in older mice for the BALB/cA strain. These observations suggest that BALB/cA has a stronger aging phenotype than C57BL/6. Indeed, of the two mouse strains, BALB/cA is known to be more susceptible to carcinogens. Thus, BALB/cA may have a less efficient DNA repair system and a more unstable rDNA region, resulting in an enhanced level of senescence.

## Materials & Methods

### Ethics statement

All experiments were approved by the Animal Experiment Ethics Committees at the Institute of Molecular and Cellular Biosciences, University of Tokyo (Exp # 0210). Experiments were performed in precise accordance with the manual provided by the Life Science Research Ethics and Safety Committee, University of Tokyo.

### Mice

Young mice (4 week-old, BALB/cAJc1 and C57BL/6JJc1) were purchased from CLEA Japan, Inc. (Tokyo, Japan) The old mice (approximately 2 year-old, BALB/cAJc1 and C57BL/6JJc1) were from this institute. For Figure S4, both the 8 week-old and 200 week-old C57BL/6JJcl mice were purchased from CLEA Japan, Inc.

### Determination of rDNA copy number in single cells

Bone marrow cells (2×10 ^7^) were isolated, washed three times with 5 ml PBS and then 1 ml of 0.005% propidium iodide (PI) (P4864; Sigma-Aldrich, St Louis, MO) was added. Each cell was sorted using a high speed cell sorter (MoFlo XDR; BECKMAN COULTER, Brea, CA) into a 96-well plate with qPCR buffer [SYBR Premis Ex TaqTM (Tli RHaseH Plus)] (RR420A; TAKARA, Tokyo, Japan) with 0.4 uM primers (Table S2) and 0.24% Nonidet P-40 (Darmanis et al., 2017). For qPCR, the plate was applied to a Thermal Cycler Dice^®^ Real Time System II (TP900; TAKARA) with the following amplification conditions; 98°C for 30 sec then 40 cycles of 95°C for 5 sec, 60°C for 30 sec. The standard curve was generated by serial dilution of DNA from Human Retinal Pigment Epithelial cells (RPE1). The rDNA copy number of RPE1 was determined by Droplet Digital PCR (ddPCR). Briefly, 5 ng RPE1 DNA was digested with HpaII (NEB, Ipswich, MA), suspended in ddPCR mixture [ddPCR Supermix (No dUTP) (1863023; Bio-Rad, Hercules, CA), target primers/probe (FAM), reference primers/probe (VIC, TaqMan™ Copy Number Reference Assay, human, RNase P, 4403326; ThermoFisher, Waltham, MA)] and applied to a X200™ Droplet Generator (1864002; Bio-Rad). Each droplet was collected into a 96-well plate [twin.tec semi-skirted 96-well plate, 951020362; Eppendorf, Enfield, CT] and detected by PCR using the following conditions; 95°C for 10 min followed by 40 cycles of 94°C for 30 sec, 60°C for 1 min and then 98°C for 10 min. The signal was detected by a QX200™ Droplet Reader and the number of positive droplets calculated using QuantaSoft™ Software (1864003; Bio-Rad). On average a RPE1 cell had 330 rDNA copies.

### Southern blot analysis to detect rDNA

For Southern blot analysis 150 ng of mouse DNA was digested with 10 units of BamHI-HF (NEB, Figure 2 and 5), NdeI (NEB, Figure 2 and 5) and SacII (NEB, Figure 5) overnight at 37°C. The digested DNA was resolved on a 0.8-1.0% agarose gel (in 1xTAE) and blotted onto a filter. The 28S and SWI5 were detected on the same filter using PCR amplified probes with specific primers (Table S3). For the psoralen crosslinking assay, 2 × 10^7^ bone marrow cells were suspended in 8 ml Opti-MEM^®^ I Reduced Serum Medium (ThermoFisher) and divided into two 6 cm-diameter dishes. A 200 µl solution of psoralen in methanol (200 µg/ml, Sigma-Aldrich) was added to each dish and only methanol to the control dish. Each of the dishes were placed on ice for 5 min and crosslinked using UV-A for 4 min (7 cm apart from the UV light). This UV exposure and psoralen addition cycle was repeated four times. Cells were then scraped and collected by centrifugation (1,800 rpm, 5 min), and the DNA isolated. A 500 ng aliquot of DNA was digested with 20 units of AflIII (NEB) overnight at 37°C and subjected to Southern blot analysis (1% agarose gel in 1xTAE, 60 V, 18 hr). The gel was then exposed to UV (4000 J/cm^2^ x 100) using a UV Stratalinker to reverse the crosslinking(42), (43), (44).

### RT qPCR

Bone marrow cells (1×10^7^) were washed with 5 ml PBS twice and the total RNA was isolated using a RNeasy Mini Kit (74104; Qiagen, Hilden, Germany). The solution was subsequently treated with DNase I (79254; Qiagen). The RNA was reverse transcribed to DNA by ReverTra Ace qPCR RT Master Mix (FSQ-201; TOYOBO, Tokyo, Japan) and the DNA solution (0.2 ng) was applied to qPCR using SYBR Premis Ex TaqTM (Tli RHaseH Plus) (RR420A; TAKARA). For normalization, housekeeping genes, Actb (actin, beta), GAPDH (glyceraldehyde-3-phosphate dehydrogenase) and B2M (beta-2 microglobulin) were also examined. The sequences of the primers (0.4 µM each) are given in Table S2. The PCR conditions were as follows; 40 cycles of 95°C for 5 sec, 60°C for 30 sec.

### rDNA sequence analysis

rDNA coding regions (18S, 5.8S and 28S) were amplified by PCR. The PCR mix included 20 ng of rDNA or 150 ng of ATP5b gene genomic DNA in a 40 ul reaction mixture (0.2 mM dNTPs, 1.5 mM MgSO_4_, 0.25 µM primers, 1× PCR Buffer for KOD-Plus-Neo, 0.8 U KOD-Plus-Neo). The sequences of the primers are listed in Table S2. The PCR cycle conditions were as follows for 18S rRNA: 94°C for 2 min, 25 cycles of 98°C for 10 sec, 60°C for 30 sec, 68°C for 90 sec, then 68°C for 15 sec; for 5.8S rRNA: 94°C for 2 min, 25 cycles of 98°C for 10 sec, 60°C for 30 sec, 68°C for 30 sec, then 68°C for 15 sec; for 28S rRNA: 94°C for 2 min, 25 cycles of 98°C for 10 sec, 68°C for 3 min, then 68°C for 15 sec; for ATP5b: 94°C for 2 min, 25 cycles of 98°C for 10 sec, 68°C for 30 sec, then 68°C for 15 sec. The PCR products were purified by Nucleospin^®^ Gel and PCR Clean-up (740609-250; TAKARA).

Purified DNA was sonicated using a Covaris M220 instrument (Covaris, Woburn, MA) to 150-200 bp long fragments. The DNA was purified using a QIA quick PCR Purification kit (QIAGEN), and the library generated with a NEB Next Ultra II DNA Library Prep Kit for Illumina (NEB). The quality of the library was checked using an Agilent 2100 Bioanalyzer (Agilent High Sensitivity DNA kit). Sequencing was performed by HiSeq 2500 (Illumina, Hiseq SR Cluster kit v4, Hiseq SBS Kit v4, 50 cycles). The sequence data was mapped on the reference sequence (GenBank BK000964) using Bowtie 2 (version 2.3.3.1), and the base frequency at each position was calculated to obtain the mutation rate (substitution, insertion and deletion) on a Galaxy platform (https://usegalaxy.org/). High-throughput sequencing data have been uploaded to NCBI Sequence Read Archive database under accession code PRJNA636244 (https://www.ncbi.nlm.nih.gov/sra).

### Yeast strains

Yeast strains expressing plasmid-borne rDNA with distinct mutations were constructed by plasmid shuffling. In brief, an rDNA depletion strain NOY986 (MATa ade2-1 ura3-1 trp1-1 leu2-3,112 his3-11,15 can1-100 *rdnΔΔ*::hisG (30) carrying a high-copy-number rDNA/URA3+ plasmid was first transformed with a rDNA/LEU2+ plasmid containing mutation at A3295(mouse A4614) or A2131(mouse A3291) within the 25S region. Strains that had lost the former URA3+ plasmid were then positively selected on SC-LEU plates containing 5-FOA.

### Yeast chronological lifespan analysis

Yeast cells were streaked on a 2%-glucose YP plate from a glycerol stock and incubated at 30°C for 3 days. A single colony was grown at 30°C overnight in 2 ml SC medium containing 2% glucose, shaking at 200 rpm. The culture was diluted with fresh 2%-glucose SC medium to an optical density of 0.1 (OD_600_ units) to give a day 0 culture of 20 ml. Starting at day 3 and every 2 days, a 100 μl aliquot of the culture was removed and diluted with sterile water to prepare a 1:10,000 dilution. The dilution was spread onto a 2%-glucose YP plate and incubated at 30°C for 3 days. The number of colony-forming units (CFU) was scored and normalized with that of the day 3 culture to establish the survival rate. All experiments were performed in biological triplicates.

## Acknowledgement

We thank Drs Yusuke Yamazumi and Tetsu Akiyama for gifting old mice and for technical advice on how to isolate bone marrow cells. We thank Drs. Yufuko Akamatsu, Mariko Sasaki, Tetsushi Iida, Chihiro Horigome and all other members of the Laboratory of Genome Regeneration for helpful discussions. This research was supported by AMED under Grant Number JP20gm1110010 to TK and T.I., and in part by grants-in-aid for Scientific Research [17H01443 to T.K.] from the Japan Society for the Promotion of Science (JSPS).

## Supporting information

### Supporting figure and table legends

**Figure S1.**
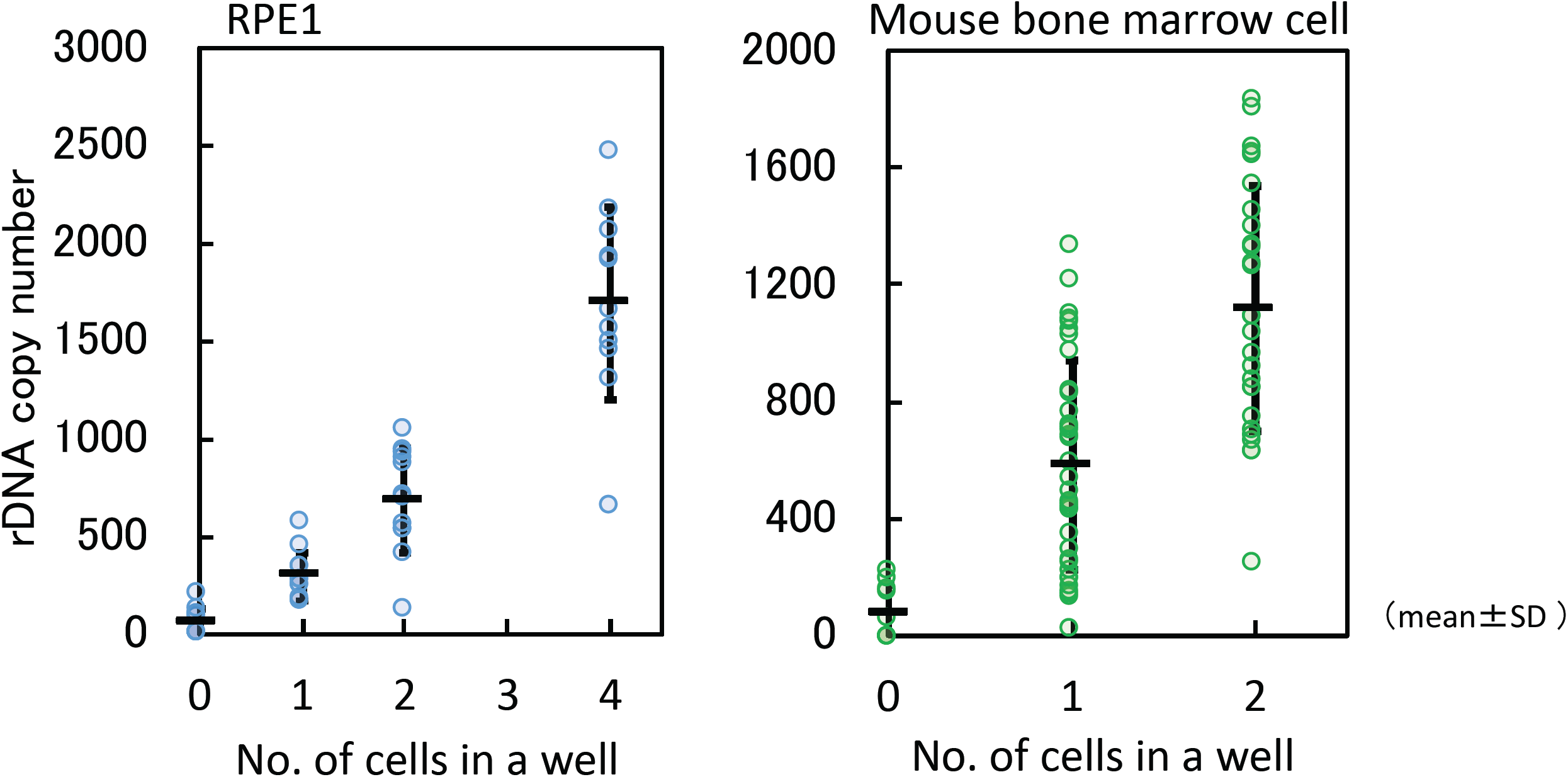
Accuracy in single cell analysis. rDNA copy numbers of one, two and four cells were measured by qPCR. rDNA copy number in RPE1 determined by digital PCR was used as the standard (Materials & Methods). In all, 12 samples for RPE1 and 36 samples for bone marrow cells were tested (dots). The horizontal bars are the average and vertical bars are S.D. Left panel: RPE1 and right panel: mouse bone marrow cell.

**Figure S2.**
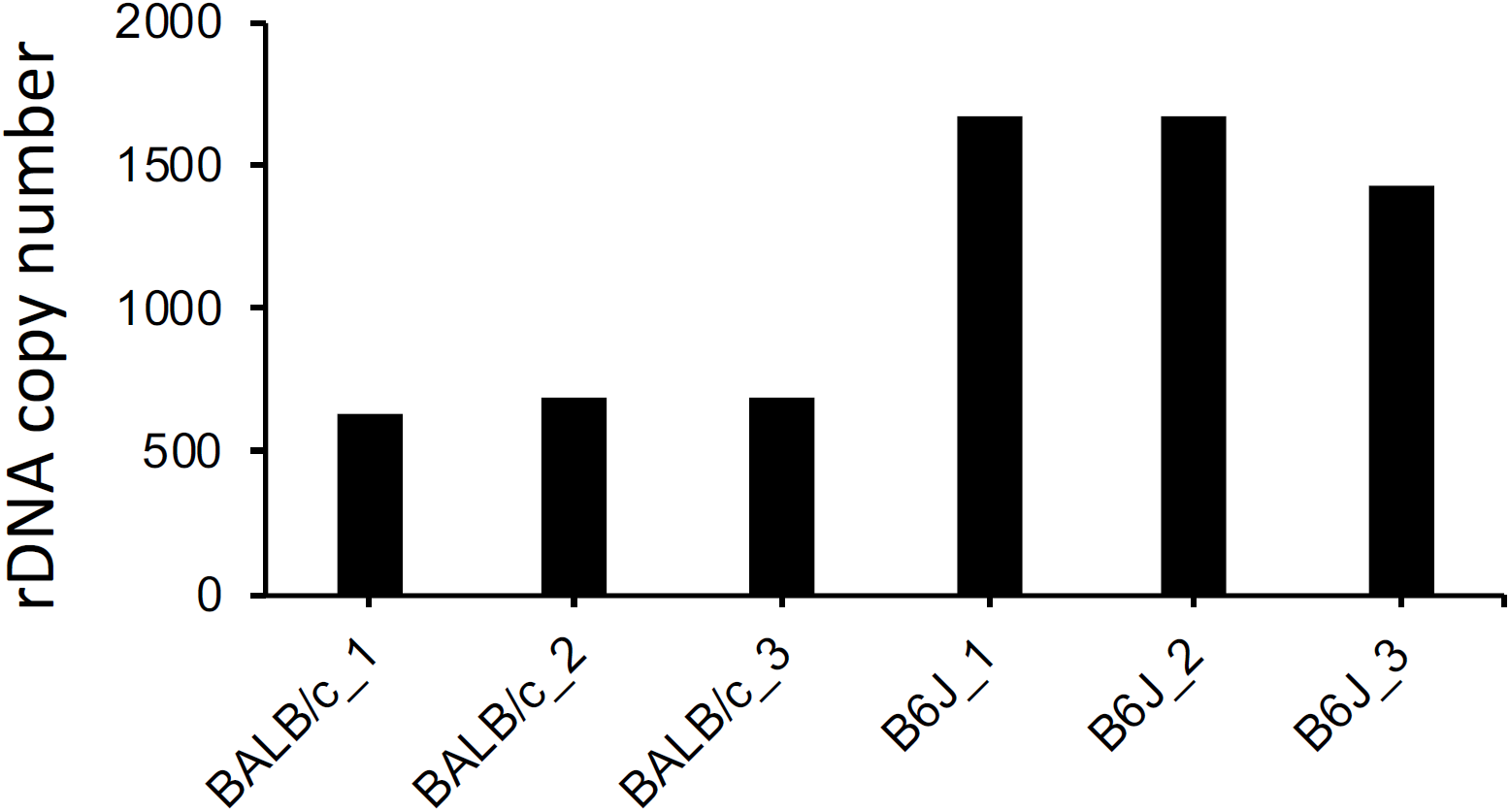
Estimated rDNA copy number from database. rDNA copy numbers of BALB/cA (BALB/c 1-3) and C57BL/6 (B6J 1-3) mice were estimated by reanalysis of publicly available whole genome sequencing data. rDNA copy number estimation by whole genome sequencing data were performed as follows. Fastq files obtained from NCBI SRA (PRJNA41995, PRJNA386034) were mapped against mouse whole genome and rDNA sequence using Bowtie2, and the fraction of rDNA reads among all mapped reads were used to calculate the copy numbers.

**Figure S3.**
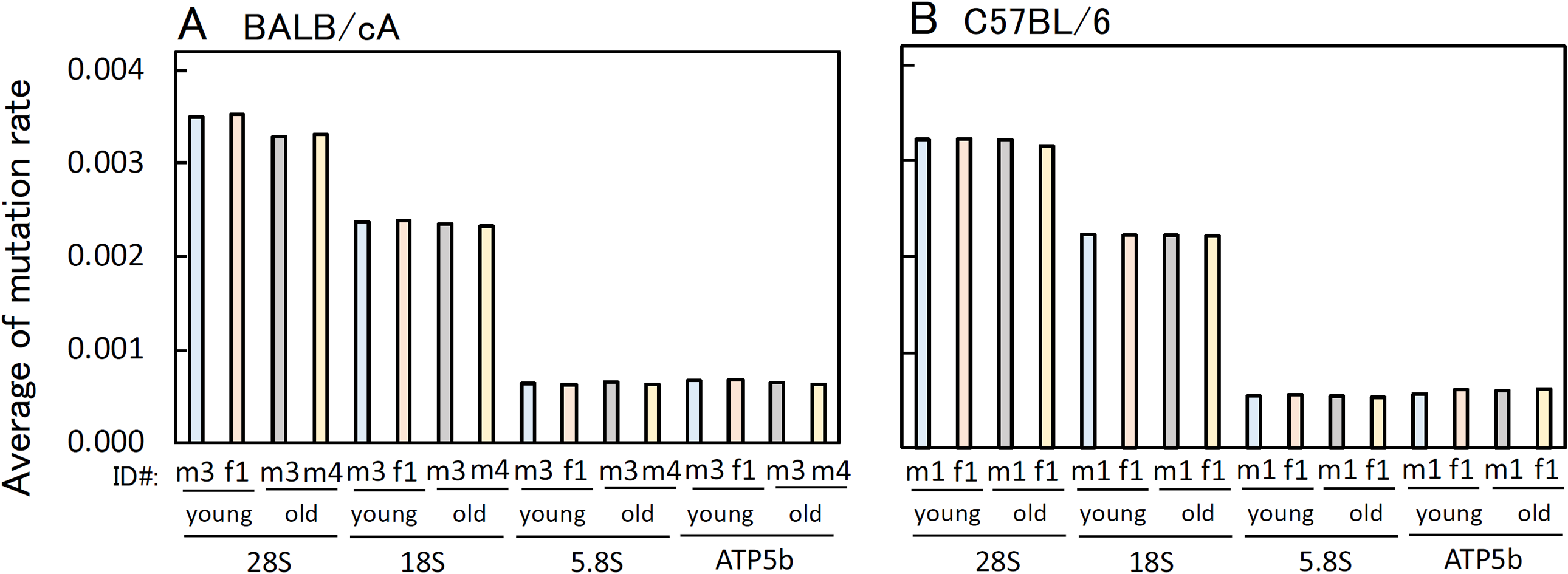
Mutation rates of young and old mice rDNA. Average mutation rates in the rDNA were calculated. (**A**) BALB/cA. young m3: young male 3, young f1: young female 1, old m3: old male 3, old m4: old male 4. The same mice were used for Figure 6. (**B**) C57BL/6. young m1; young male 1, young f1: young female 1, old m1: old male 1, old f1: old female 1. The same mice were used for Figure 6. Mutation rates that are more than 0.9 were not used because they are different from the reference sequences.

**Figure S4.**
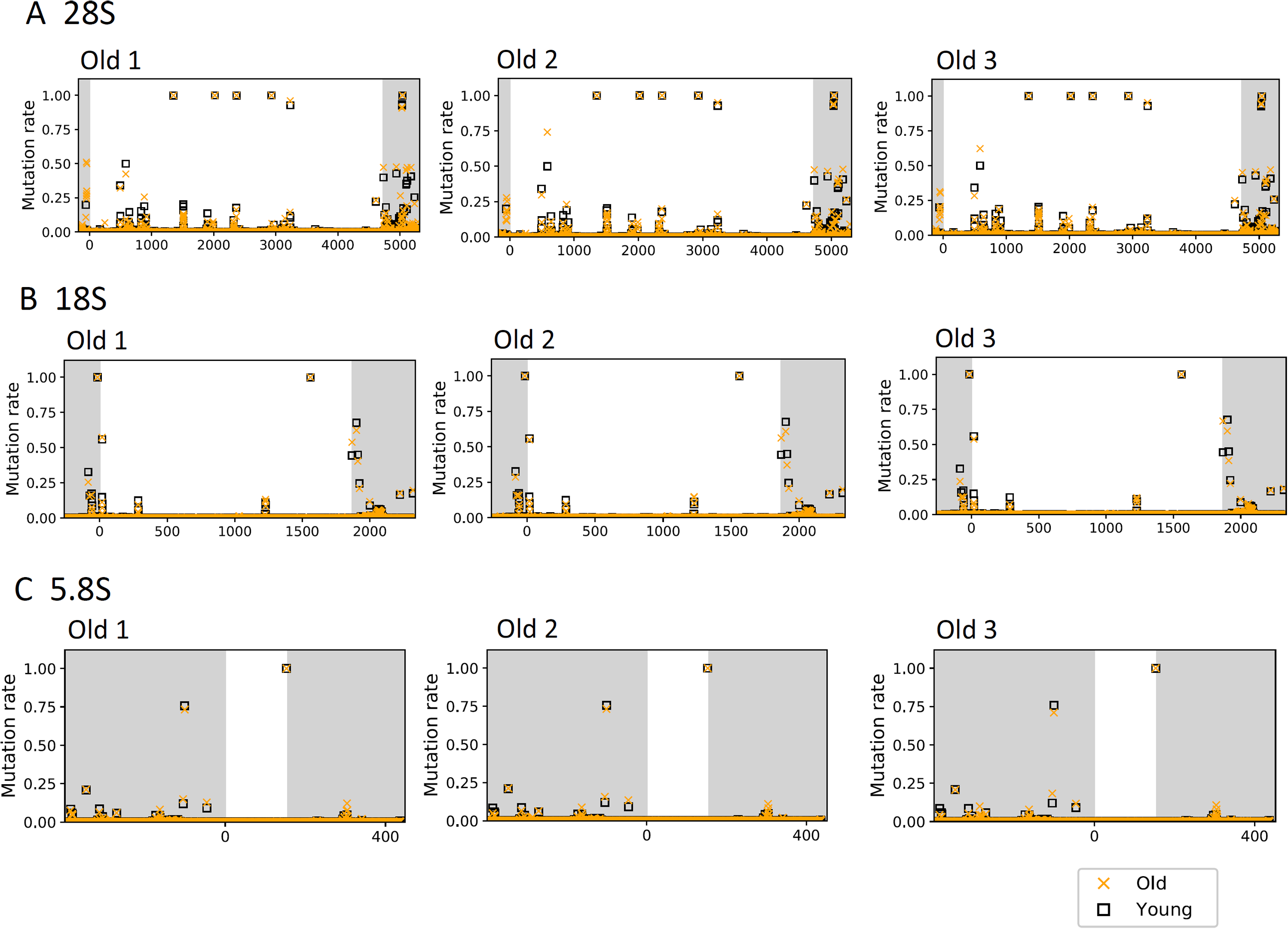
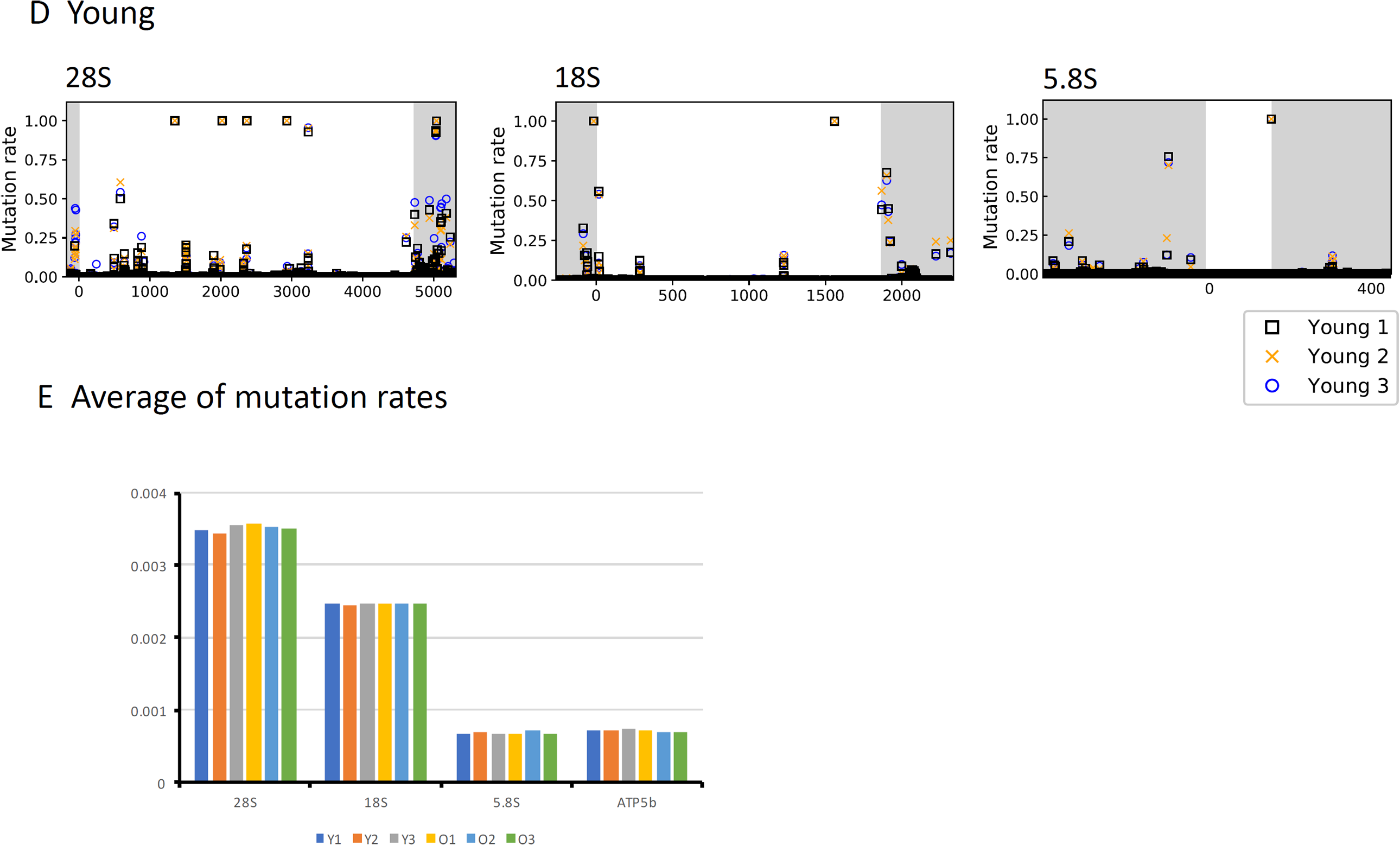
rDNA sequence variation in young and old C57BL/6 mice. rDNA sequences from three old (old 1-old 3) and three young (young1-young3) mice (C57BL/6) were determined and compared to the reference sequence. The calculated mutation rates at each nucleotide position was determined as for Figure 6. (**ABC**) Old 1-3 mice were compared with young 1 mouse. (**A**) 28S rRNA gene, (**B**) 18S rRNA gene and (**C**) 5.8S rRNA gene, (**D**) Three young mice and (**E**) Average of mutation rates. Non-coding regions are shadowed. Ave. is the average mutation rate. Mutation rate is the difference from the reference sequence as for Figure S3.

**Table S1.** Sequence variation at BamHI recognition sequences in young and old mice. Mutation rates of BamHI sites in young and old BALB/cA mice. (Upper panel) original BamHI sequence and its mutated version. (Lower panel) Mutation rates of BamHI sites. Ave. is the average sum of the mutation rates.

| Position in<br>28S | Mutation rate |  |  |  |
| --- | --- | --- | --- | --- |
|  | young |  | old |  |
|  | male 3 | female 1 | male 3 | male 4 |
| G5050 | 0.00 | 0.00 | 0.00 | 0.00 |
| G5051 | 0.00 | 0.00 | 0.00 | 0.00 |
| A5052 | 0.23 | 0.27 | 0.35 | 0.32 |
| T5053 | 0.00 | 0.00 | 0.19 | 0.16 |
| C5054 | 0.00 | 0.00 | 0.19 | 0.16 |
| C5055 | 0.00 | 0.00 | 0.00 | 0.00 |

| Position in<br>28S | Mutation rate |  |  |  |
| --- | --- | --- | --- | --- |
|  | young |  | old |  |
|  | male 1 | female 1 | male 1 | female 1 |
| G5050 | 0.00 | 0.00 | 0.00 | 0.00 |
| G5051 | 0.00 | 0.00 | 0.00 | 0.00 |
| A5052 | 0.22 | 0.25 | 0.21 | 0.21 |
| T5053 | 0.09 | 0.11 | 0.09 | 0.08 |
| C5054 | 0.08 | 0.11 | 0.09 | 0.08 |
| C5055 | 0.00 | 0.00 | 0.00 | 0.00 |

Ave: 0.44
Ave: 0.38

**Table S2.**
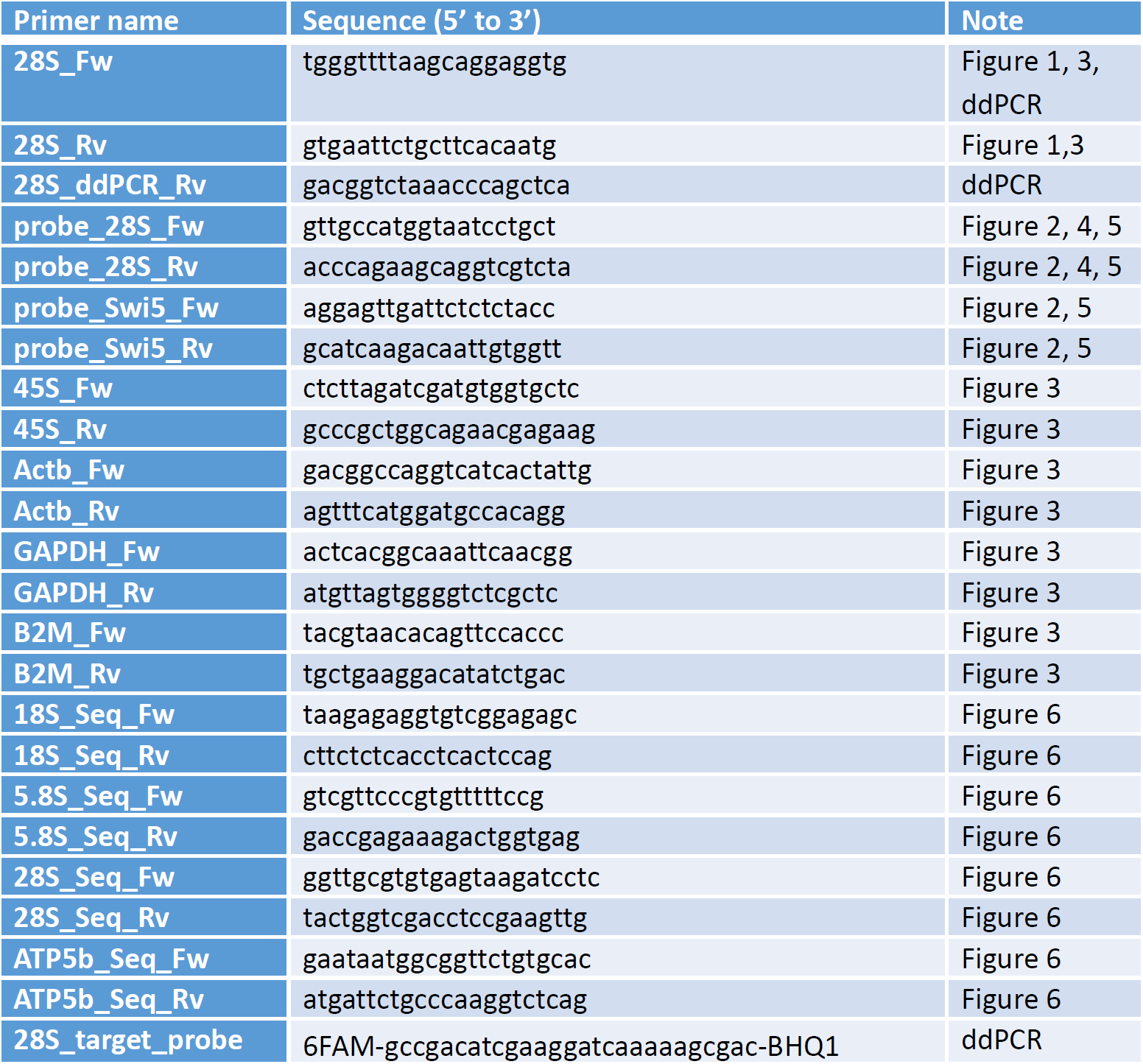
Primer list.

